# Development of a metatranscriptomic analysis method for multiple intestinal sites and its application to the common marmoset intestine

**DOI:** 10.1101/2021.10.12.464166

**Authors:** Mika Uehara, Takashi Inoue, Minori Kominato, Sumitaka Hase, Erika Sasaki, Atsushi Toyoda, Yasubumi Sakakibara

## Abstract

**Background:** The intestinal microbiome is closely related to host health, and metatranscriptomic analysis can assess the functional activity of microbiomes by quantifying the bacterial gene expression level, which helps to elucidate the interaction between the microbiome and the environment. However, functional changes in the microbiome along the host intestinal tract remain unknown, and previous analytical methods have limitations, such as potentially overlooking unknown genes due to dependence on existing databases and being unable to take full advantage of metatranscriptome to reveal the functional change among multiple environments.

**Result:** To close these gaps, we develop a novel method that integrates metagenome and metatranscriptome to analyze the functional activity of microbiomes between intestinal sites. This method reconstructs a reference metagenomic sequence across multiple intestinal sites, allowing the gene expression levels of microbiome including unknown bacterial genes to be compared between multiple sites. As a result of applying this method to metatranscriptomic analysis in the intestinal tract of common marmoset, the reconstructed metagenome covered most of the expressed genes and it revealed that the changes in bacterial gene expressions among the caecum, transverse colon, and faeces were more dynamic and sensitive to environmental shifts than its abundance. In typical, the coenzyme synthesis gene and antibacterial resistance gene were more highly expressed in the caecum and transverse colon than in faeces, while there was no significant change in abundance of these genes.

**Conclusion:** Our findings demonstrate that an analytical method that integrates metagenome and metatranscriptome in multiple intestinal sites captures functional changes in the microbiomes at the gene resolution level.

## Introduction

In the past few decades, many sequence-based analyses have attempted to elucidate the relationships between microbiomes and environments such as the ocean, soil, and digestive tract. These studies have traditionally focused on profiling membership through amplicon sequencing of the 16S rRNA gene. Recently, whole metagenomic sequencing methods, which enable comprehensive capture of microbial genomes to reconstruct database-independent metagenome sequences and reveal potential microbial genes and the community taxonomic abundance profiles, have become more widely used due to recent advances in sequencing throughput and analytical methods. For instance, in a large-scale metagenomic analysis spanning human body parts—the oral cavity, skin, faeces, and vagina—154,723 microbial genomes were reconstructed, 77% of which were unknown genomes not found in public repositories (1). Additionally, a study on the cow rumen microbiome reported 913 microbial genomes, and these reconstructed genomes improved the metagenomic read classification by sevenfold (2). Other studies have shown that microbial genes detected on reconstructed metagenomic sequencing play an important role in pathology of rheumatoid arthritis (3). Although these metagenomic studies have revealed many insights into a wide variety of microbiomes by finding new bacterial genomes and potential genes and emphasized the importance of reconstructing bacterial genomes, these approaches show only the presence of microbiome members and their genes and cannot indicate whether they are active members of the microbiome or how the bacteria actually interact with the environment. As a way to solve these problems, metatranscriptomic analysis records expressed transcripts within a microbiome to obtain deeper insight into how bacterial communities respond to environmental conditions. A study that included both metatranscriptomic and metagenomic analyses in patients with inflammatory bowel disease (IBD) highlighted the metabolic pathways characteristic of the disease and revealed whether metagenomically abundant bacteria were inactive or dormant in the intestine (4). In a human faecal microbiome study with both metagenomic and metatranscriptomic analysis, the metatranscriptome was more dynamic than the metagenome, and there was a discrepancy between bacterial abundance and transcriptional activity (5). As such, finding microbial gene expression signatures can be crucial to understanding the mechanisms behind microbe-environment interactions.

Metatranscriptomic analysis utilizes three main approaches to quantify bacterial transcripts, each with corresponding drawbacks. The first is the read-based approach used in the pipelines such as HUMAnN2 (6) and SAMSA2(7), which assess the activity of each protein family and pathway by aligning reads derived from metatranscriptomic library preparations with protein databases such as RefSeq (8) and pathway databases such as KEGG (9) and MetaCyc (10), respectively. This method is simple and often used but may missed many previously unknown genes that are not annotated in the databases.

The second approach is de novo assembly of RNA reads with programs such as Trinity (11) and SOAPdenovo-Trans (12). In this method, the transcript is reconstructed from the RNA short reads by de novo assembly, and the expression level is quantified by aligning the RNA read with this transcript. This method does not rely on the databases, whereas these assemblers are designed for a single organism and have not been shown to be effective in accurately assembling transcripts from a complex community (13).

The third approach performs metatranscriptomic analysis based on de novo assembly of metagenomic data. Gene expression is quantified by aligning RNA reads with the predicted genes for contigs obtained by assembling corresponding metagenomic DNA reads, which requires simultaneous sampling of the metagenome and metatranscriptome from the same sample. This approach is powerful enough to discover and focus on unknown genes and is therefore adopted in this study as well. When applied to the analysis of microbiome in multiple environments, the difficulty of this approach is to identify the same gene across samples because the assembled genomic sequence varies from base to base depending on samples. This limitation prevents comparison of the bacterial gene expression level between different environments such as multiple intestinal sites. In previous studies that attempted to gain insight into newly discovered genes, unannotated genes predicted on the reconstructed genome were clustered by sequence similarity and the gene activity was assessed by summing the expression levels of genes within the same cluster (14). However, in this approach, all similar genes encoded in multiple bacterial species are combined into a single one, and it is thus still not possible to quantify the expression level of each gene in each bacterial species.

The intestinal tract regulates highly complex physiological processes while interacting with a dense and diverse microbial population. Most studies use faecal sample on the assumption that faeces reflect the condition of the microbiome inside the intestinal tract (1) (3) (4) (5). Since the function of the intestinal tract varies from site to site, and there are differences in the physicochemical environment, such as nutrients, oxygen, and pH, the microbiome may differ in response to changes in the environment (15) (16). Indeed, due to these environmental shifts, some studies have reported that the composition of the microbiome varies depending on the intestinal sites in model animals, such as mice and pigs (17) (18) (19) (20). However, these studies have shown only differences in the microbial members in the intestinal tract, and it is still unclear how the microbial function varies along the intestinal tract. Moreover, for the aim of applying it to the interrelationship between humans and microbiomes, we need to study using an animal model that are more anatomically and pharmacologically resemble to the human. The common marmoset is a small new world primate that is considered a useful model in preclinical studies due to its common physiological and anatomical characteristic with those of human (21).

In the present study, we aim to clarify the changes in microbial abundance and gene expression due to environmental gradients among the caecum, transverse colon, and faeces. To accurately perform this investigation, it is necessary to overcome the discrepancy between the microbes existing in the environment and those registered in the databases such as COG and KEGG Orthology (KO) database (1) (2). Therefore, we developed an integrated metagenomic and metatranscriptomic method for analysis of the functional changes in microbiomes across multiple intestinal sites and then applied this method to the investigation of common marmoset intestine.

## Results

### Overview of the proposed analytical method that integrates metagenome and metatranscriptome to analyze the functional activity of microbiomes between intestinal sites

After assembly and scaffolding of the metagenomic reads, the proposed analytical method used a strategy to reconstruct the common reference metagenome, including those of unknown bacteria, by merging the scaffolding between samples; accordingly, the expression levels of all bacterial genes can be quantified by integrating this reconstructed reference metagenome with metatranscriptome data. The overview of the proposed analytical method is illustrated in Fig. 1. Using this method, we compared the microbial gene expression levels among three sites—the caecum, transverse colon, and faeces. These sites were selected as locations equivalent to the proximal, middle, and distal position of the colon, where the most bacteria are located (22). In addition, we compared the corresponding microbial compositions among humans, mice, rats and marmosets to evaluate the suitability of common marmosets as an animal model in microbiological studies.

**Fig. 1.**
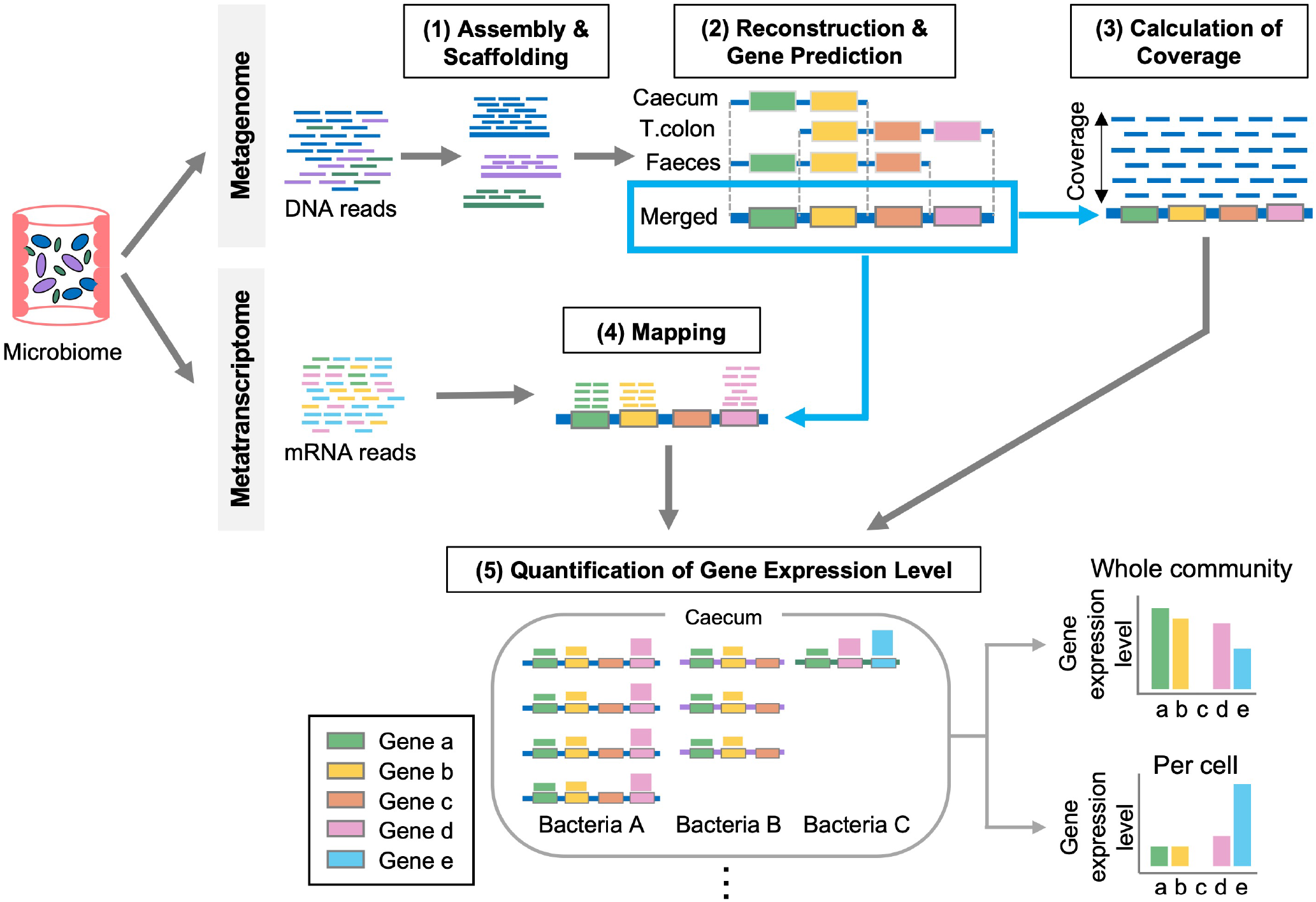
Overview of the proposed analytical method. This method integrates metagenome and metatranscriptome to analyze the functional activity of microbiomes between intestinal sites as follows. Samples for the metagenome and the metatranscriptome are taken simultaneously. (1) Assembly with DNA reads generates contigs and scaffolds at each site. (2) Bacterial metagenomes are reconstructed by merging scaffolds across all sites. Gene-coding regions are predicted on the reconstructed metagenome. (T.colon represents the transverse colon.) (3) DNA reads are mapped to the reconstructed metagenome to calculate relative abundance. (4) mRNA reads are aligned to the reconstructed metagenome and, mapped reads are quantified for each gene. (5) Gene expression levels are calculated at whole community. Gene expression levels per cell are calculated by normalizing with gene abundance.

### Metagenome merging-reconstruction improves assembly contiguity, transcript mapping rate and identification of same genes between sites

A total length of 306 Mb and 395 Mb reference metagenomes consisting of 32,244 and 39,905 scaffolds were reconstructed by merging from three intestinal sites for individuals 1 and 2, respectively. We compared scaffold length before and after merging scaffolds from three sites by a generalized score N-statistic, which is an extension of N50. Scores from N10 to N100 were plotted at 10 intervals in Fig. 2A. The genome assembly of each intestinal site fell well below in comparison to the merged one. This implied that merging improved the assembly contiguity, which means that the scaffolds of three sites complemented each other to reconstruct a longer genome. Next, a total of 246,980 and 320,613 genes were detected in the reconstructed metagenomes for individuals 1 and 2, respectively. Of the genes detected in individuals 1 and 2, 63,331 and 88,575 (26% and 28%) genes were not present in the COG database, and 112,790 and 152,845 (46% and 48%) genes were not present in the KEGG database (Fig. 2B; Table S1). Thus, a large number of novel genes not included in the public database were detected on the reconstructed metagenomes.

**Fig. 2.**
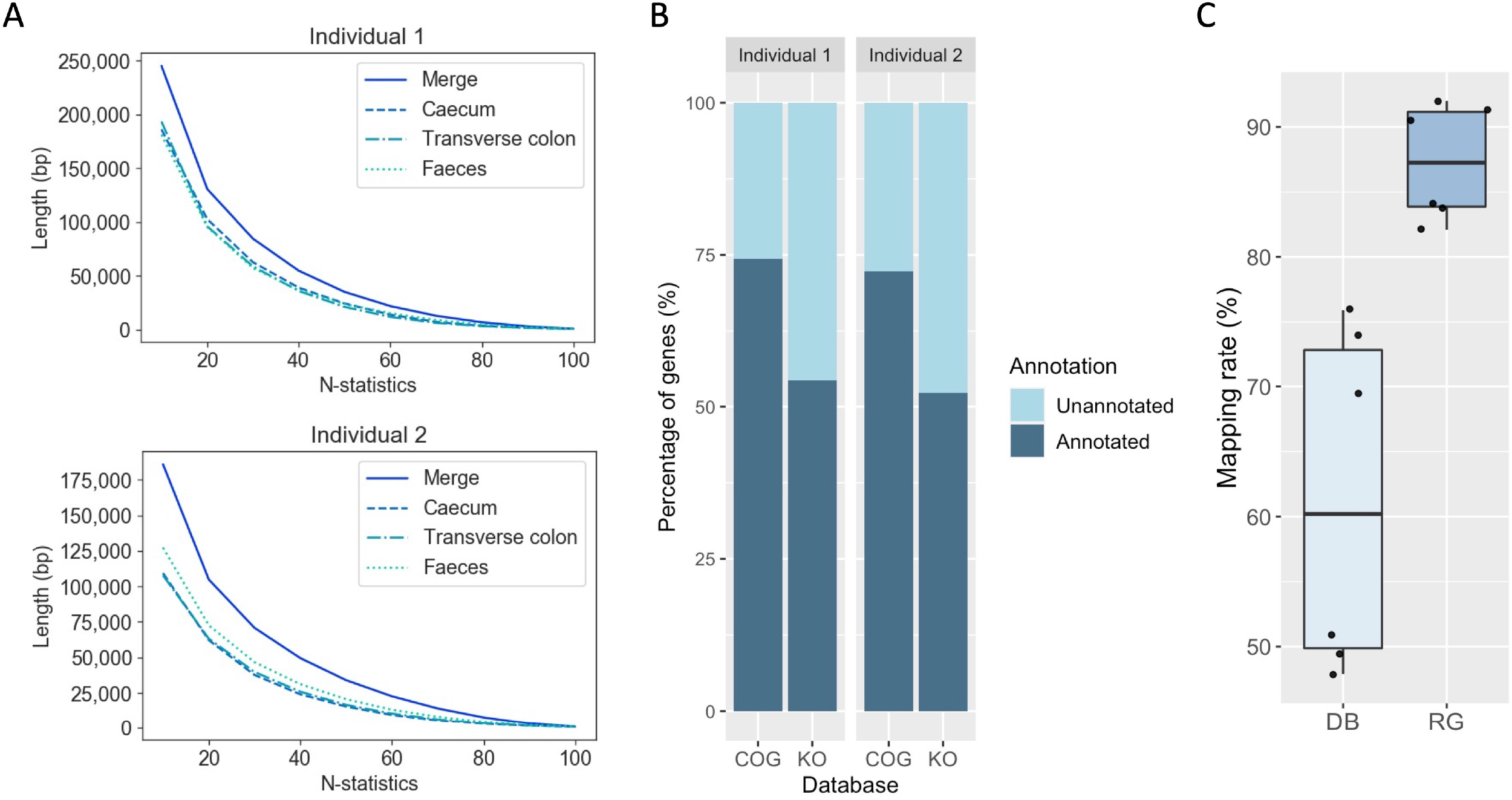
Reconstruction of a merged metagenome improved the assembly contiguity, gene detection and read mapping rate. (A) The plots of N-statistics to measure the assembly contiguity reconstructed from three intestinal sites and of the merged metagenome in individuals 1 and 2. We computed the N-statistics from N10 to N100 at 10 intervals, which is an extension of N50 measure to evaluate the assembly contiguity. (B) Percentage of functionally annotated genes in the reconstructed genomes. Approximately 63,331 and 88,575 genes are not present in the COG database, and 112,790 and 152,845 genes are not present in the KO database. (C) Mapping rate of microbial mRNA reads to the database and the reference metagenome (DB = database, RG = reconstructed reference metagenome). This boxplot represents the mapping rates of mRNA reads from the caecum, transverse colon, and faeces in individuals 1 and 2, respectively.

To quantify the gene expression level, we first mapped the mRNA reads to all complete bacterial, archaeal, and viral genomes in the RefSeq database (8). Only 21–52% of mRNA reads could be assigned to the known genomes (Fig. 2C). This result confirmed that information to understand the microbiome activity was limited if relying solely on the genomes registered in the public database. We therefore mapped the mRNA reads to the reference metagenomes reconstructed in this study. The mapping rate to the reconstructed metagenomes increased to 82–90% (Fig. 2C). The reconstructed metagenomes covered most of the expressed genes (Table S2) and allowed us to map 2-4 folds more reads than the database. These results underscored the importance of database-independent analytical methods, especially in metatranscriptomic analysis to quantify microbial gene expression levels.

In addition, we verified that the gene annotations were retained before and after merging by examining the percentage of genes common to three sites that matched the corresponding gene in the reconstructed metagenome. We found that 96.9% and 96.6% of the genes common to the three sites were identical to those of the merged metagenome, in individuals 1 and 2, respectively (Allowed for a 3-base mismatch; Supplementary Note 5; Table S3). The reference metagenome reconstructed by merging thus achieved the high accuracy to identify the same gene between three intestinal sites.

### Functional annotation for unknown genes with metatranscriptomic profiles

To address unknown genes that were not annotated by the databases, we generated a gene catalogue from the reconstructed metagenomic sequences by grouping into gene clusters and performing a co-variation analysis. Of the unknown genes detected in two individuals, 50,509 expressed genes were grouped into 24,725 gene clusters by protein sequence similarity (Table S4). In addition, we performed a co-variation analysis that estimated the function of those unknown gene clusters (23), incorporating a bivariate spatial relevance (24) between multiple intestinal sites. We first evaluated the rationale of this co-variation analysis that pairs of genes with similar expression profiles were associated with a common metabolic process (Supplementary Note 6; Table S5). As a result of benchmarking the co-variation analysis using the gene expression level at whole community and per cell, the area under the curves (AUCs) were 0.830 and 0.729, respectively (Fig. 3A, B, and C). This co-variation analysis was then applied to the unknown gene clusters and showed that the function of many unknown gene could be involved in the xenobiotics biodegradation, energy metabolism, nucleotide metabolism, signal transduction, and digestive system (Fig. 3D; Supplementary Note 6 and 7; Table S6), which also suggests that current database-dependent analytical methods may underestimate these functions in our data. Thus, the co-variation analysis incorporating a bivariate spatial relevance, combined with metatranscriptomic analysis, provided an accurate functional interpretation of unknown genes on the reconstructed metagenome.

**Fig. 3.**
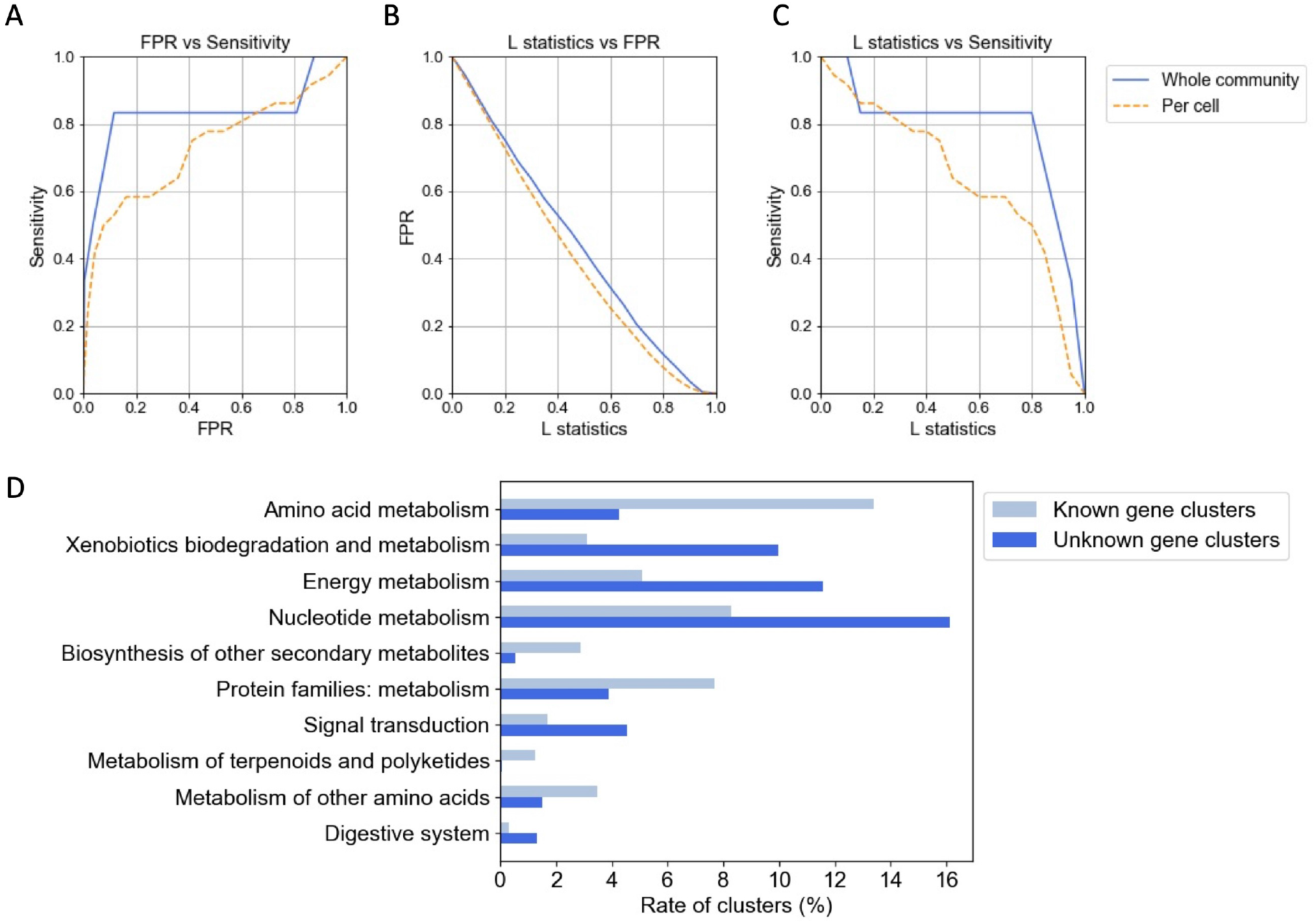
Rationale of the co-variation analysis and the resulting molecular functions associated with unknown genes. The co-variation analysis incorporating a bivariate spatial relevance was performed to associate unknown genes with molecular functions. Evaluation of the co-variation analysis using the gene expression profile at whole community and per cell: (A) Receiver operating characteristic (ROC) curves, (B) false positive rate (FPR) and (C) sensitivity along the L statistic value to associate known gene cluster pairs. True positives were defined as pairs of covariant genes with a common KEGG reaction definition. (D) The functions of unknown gene clusters associated by co-variation analysis using the gene expression profile at whole community, and the functions of known gene clusters. Only functions enriched in either unknown or known gene clusters are shown (Fisher’s exact test with p-value < 0.01 adjusted by the Benjamini-Hochberg method; Supplementary Note 6; Table S6). The L statistic value that ensured a false positive rate <5% in the benchmark was used as the threshold (Supplementary Note 6).

### Spatial variance in microbial gene expression at whole community and individual cell levels

The functional activity of the microbiome in the caecum, transverse colon and faeces was assessed using the gene expression level at both the whole community and per cell levels. The gene expression level at whole community provides functional profiling of the entire microbiome but is affected by the abundance of bacteria; on the other hand, the gene expression level per cell provides the gene activity for each bacteria, even for minority bacteria.

We extracted the biochemical functions whose expression levels varied significantly between the intestinal sites. The top 50 KOs (KEGG Orthologies) with highest expression differences between sites are shown in Fig. 4. KO identifiers, K02041: phosphonate transport system ATP-binding protein and K18910: D-psicose/D-tagatose/L-ribulose 3-epimerase were detected as differentially expressed between the caecum and transverse colon (Fig. 4A). The differentially expressed between the caecum and faeces were KO identifiers, K08260: encoding adenosylcobinamide hydrolase, K03486: GntR family transcriptional regulator, trehalose operon transcriptional repressor, and K00332: NADH-quinone oxidoreductase subunit C (Fig. 4B). In addition to the differentially expressed gene between the caecum and faeces, KO identifier K19075: CRISPR-associated protein Cst2 was differentially expressed between the transverse colon and faeces (Fig. 4C). Here, we focus on genes involved in well-studied metabolism processes. The genes that are more highly expressed at whole community, in the caecum and transverse colon than faeces were genes involved in biosynthesis of the vitamin B_12_ (*cbiZ* and *pduX*), vitamin K_2_ (*mqnE*), vitamin B_7_ (*bioD*), and vitamin B_6_ (*pdxH*), and antibiotic resistance genes (*arnA* and *arnB*) (Fig. 4B and C). The gene *cbiZ*, which salvages cobinamide (Cbi), a precursor of AdoCbl, has its roots in the archaea and was acquired by several bacterial strains via horizontal gene transfer (25). This gene is required for bacterial growth on acetate (26). The detection of *pduX* as a differentially expressed gene along with *cbiZ* is consistent with a previous study showing that *pduX* is required for the *cbiZ* mediated pathway (27). Two genes *arnA* and *arnB* are known to result in resistance to antibiotics by modifying of the outer membrane by lipopolysaccharide. This modification is regulated by the PmrA/PmrB two-component regulatory system, which is switched on by low pH (28).

**Fig. 4.**
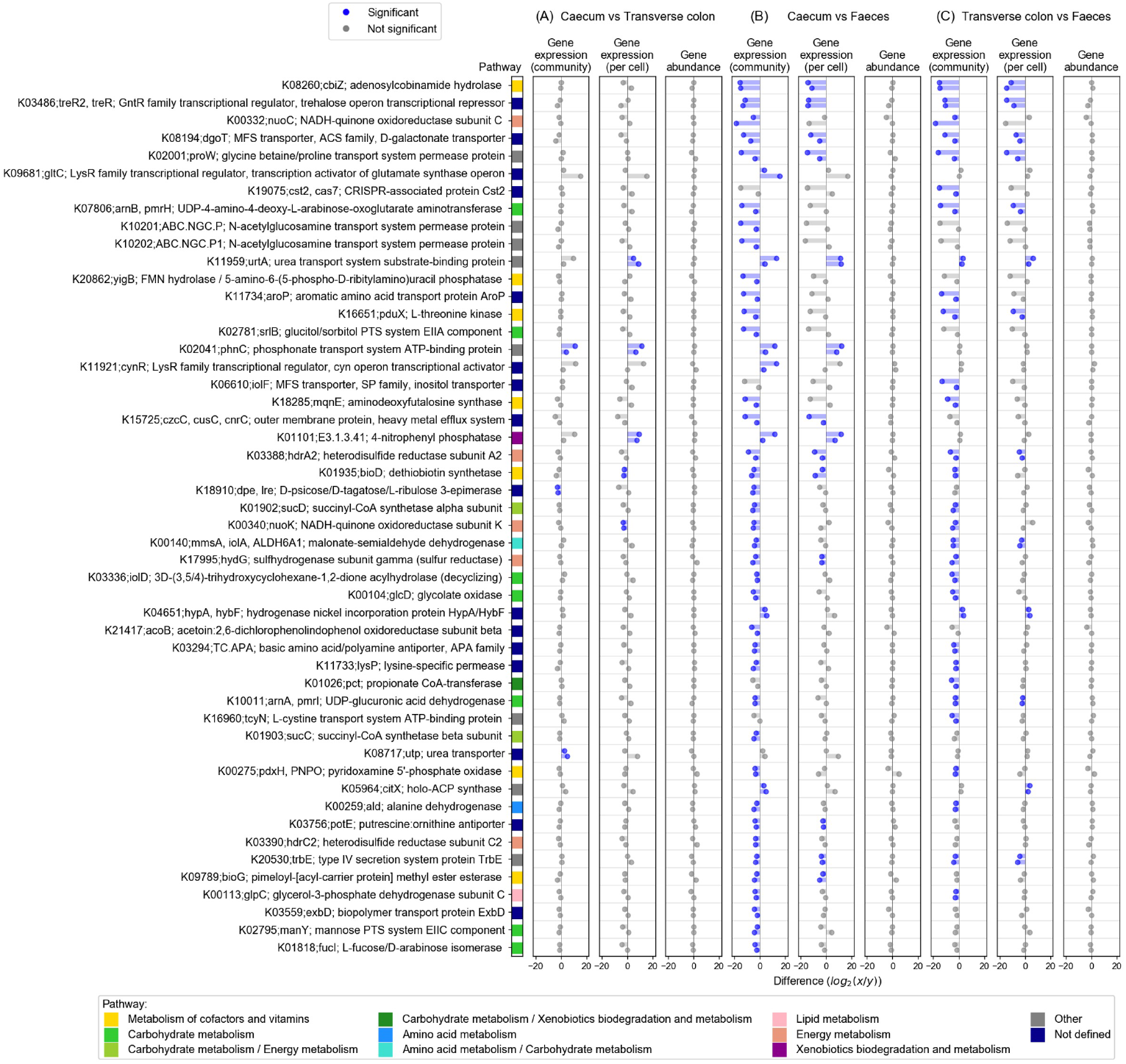
Significant KEGG Orthology in differential gene expression between the caecum, transverse colon and faeces. The top 50 KOs with highest differential gene expression in the whole community are shown, along with gene expression levels per cell and gene abundance. (A) Differential gene expression between the caecum and transverse colon; (B) Differential gene expression between the caecum and faeces; and (C) Differential gene expression between the transverse colon and faeces. The difference in gene abundance / expression levels between at whole community and per cell is displayed using log_2_-transformed values. In each KO, the upper bar represents individual 1 and the lower bar represents individual 2. The difference was considered and denoted as “significant” if the difference changed in the same direction by more than 2-fold in both of the two individuals.

These differentially expressed genes are related to sugar utilization in the intestinal tract (Fig. 5). The genes with differential expression at whole community between the caecum and faeces were the genes involved in the utilization of sorbitol (*srlB*), mannose (*manY*), and L-fucose (*fucI*) (Fig. 5(1)). This result likely reflects the utilization of sugars that were not absorbed in the small intestine by the microbiome (29). Fermentation of these sugars by the caecal microbiome produces short-chain fatty acids (SCFAs) (30), which increases the concentration of SCFAs in the colon, but it decreases in faeces due to its absorption at the colon (31). Acetic acid accounts for approximately 60% of SCFAs in the colon (32), and therefore this change in the concentration of SCFAs along the colon explains the changes in expression of *cbiZ* (Fig. 5(2)), which is essential for bacterial growth on acetate (26). Similarly, the decrease in the concentration of SCFAs from the caecum to the descending colon was accompanied by an increase in pH, which is consistent with the changes in the expression of the antibiotic resistance genes *arnA* and *arnB*, which are switched on at low pH (28) (Fig. 5(3)). Thus, many of these genes differentially expressed between intestinal sites are involved in the SCFAs produced by microbial sugar metabolism. Since these typical genes are obviously encoded in multiple bacterial species, we picked up the L-fucose metabolic gene (*fucI*) and investigated which bacteria caused the differential expression of this gene. On the reconstructed reference metagenome, 30 loci encoding *fucI* were detected, each representing one bacterial species (Fig. 6). As a result of this analysis, many bacteria belonging to the *Firmicutes* phylum contributed to the expression level of the *fucI* gene at whole community, and the scaffold ID S123510 belonging to *Megamonas* genus was a particularly important contributor in individual 1. On the other hand, the scaffold ID S127859 belonging to *Akkermansia* genus, a well-known SCFA-producing bacteria (33), was most abundant in terms of gene abundance of *fucI* in individual 1, although its expression level per cell was low. This finding demonstrates the power of our method of integrating metagenome and metatranscriptome to enable analysis at the gene resolution level.

**Fig. 5.**
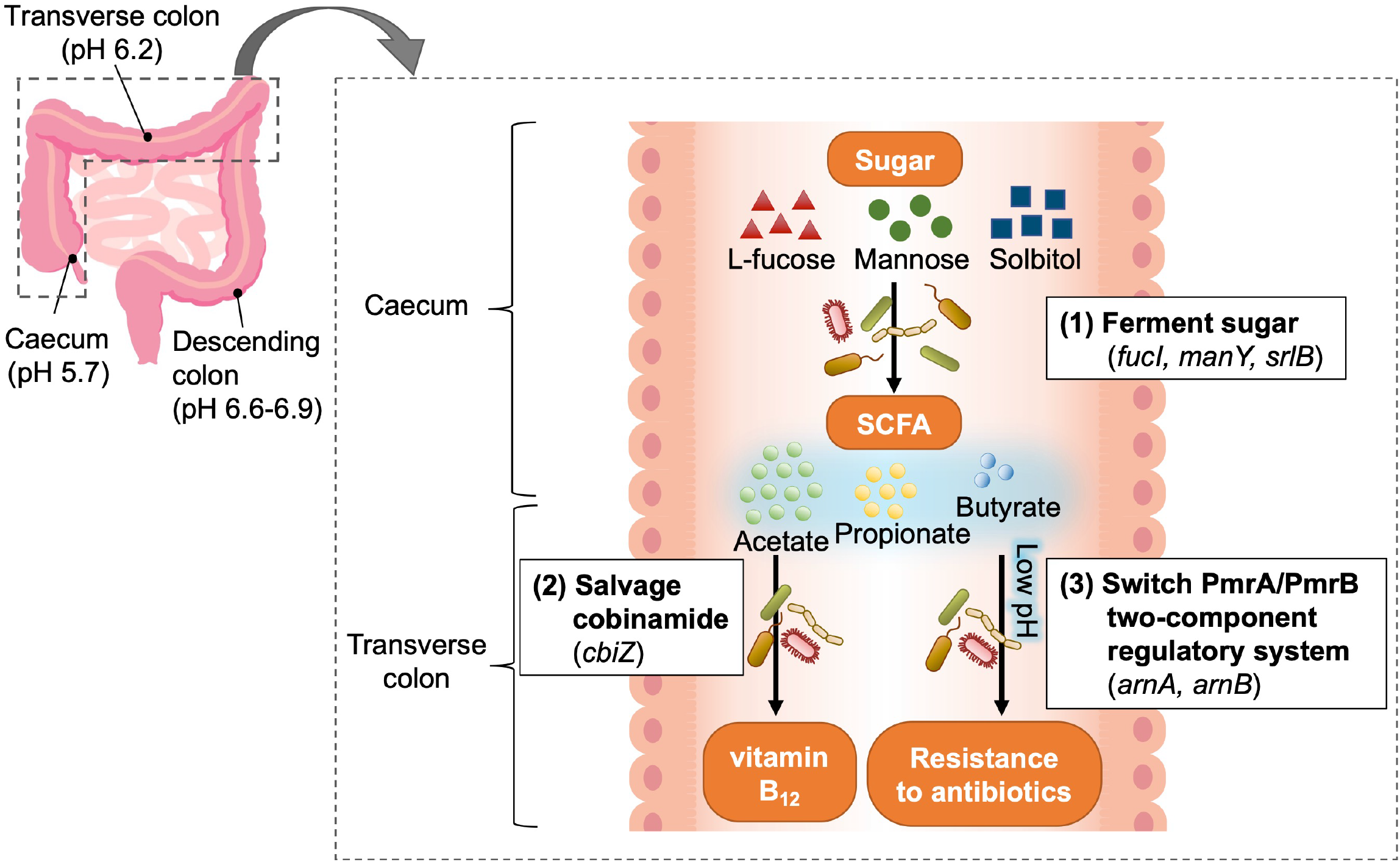
Functional activities in the microbiome along the host intestinal tract. In relation to Fig. 4, the functional shifts of the microbiome along the intestinal tract estimated from the differentially expressed genes between sites were as follows: (1) Sugars that are not absorbed in the small intestine are fermented by the caecal microbiome to produce SCFAs (29) (30). Since SCFAs are absorbed in the large intestine, the concentration of SCFAs gradually decreases from the caecum to the descending colon (31). (2) The growth of bacteria under acetate (26), which is abundant in SCFAs (32), requires *cbiZ* in the vitamin B_12_ biosynthetic pathway, and the production of SCFAs makes this gene more active in the caecum and transverse colon than faeces. (3) Similarly, a decrease in pH with increasing SCFA concentration switches on the antibiotic resistance genes *arnA* and *arnB* (28).

**Fig. 6.**
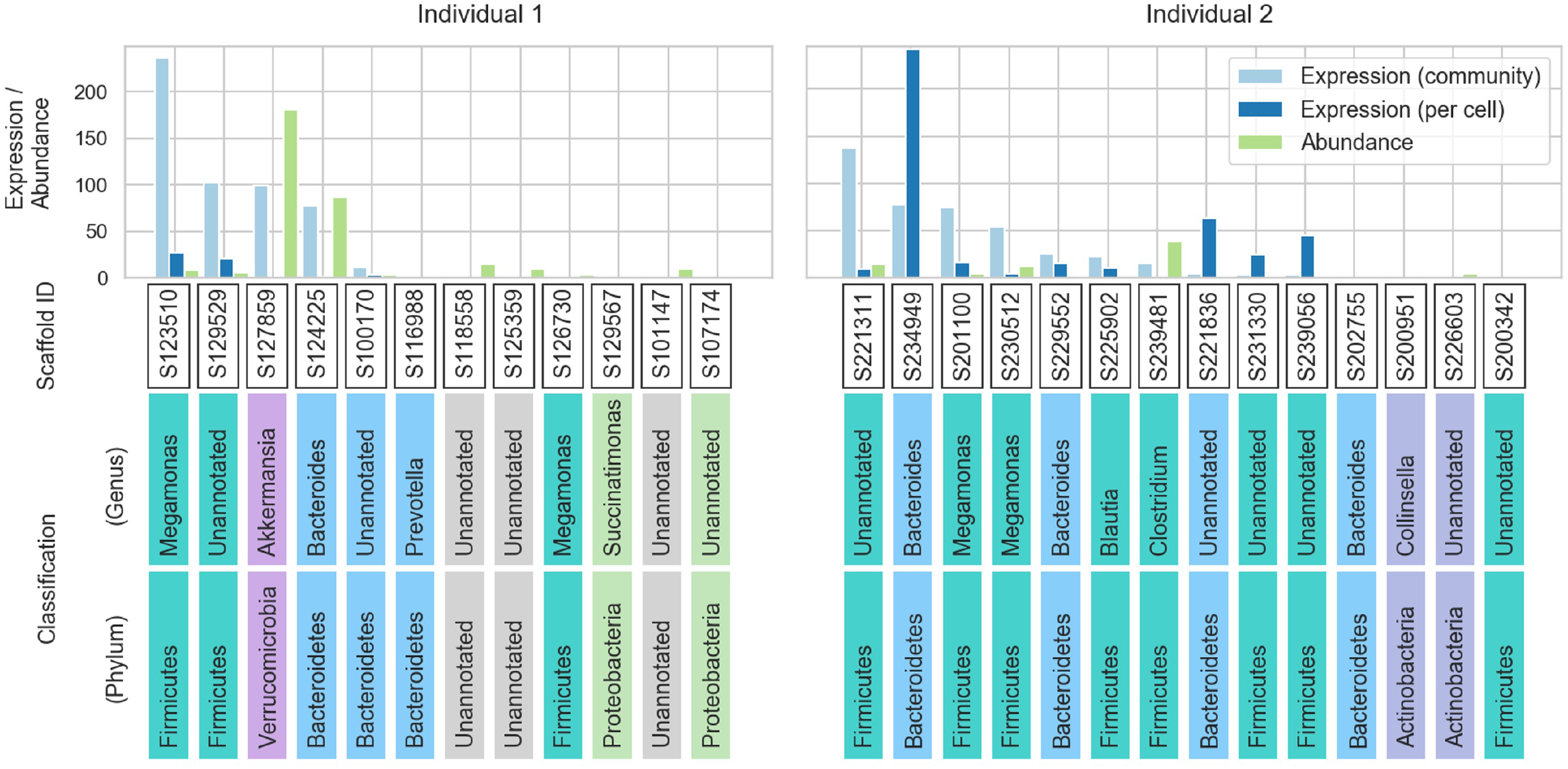
Thirty scaffolds that encode L-fucose isomerase gene (*fucI*), each representing one bacterial species. Bar plot shows the relative gene abundance and gene expression level at whole community and per cell of *fucI* on 30 scaffolds in individual 1 and 2. Each scaffold ID and its taxonomic classification is shown at the bottom, and colored by phylum level classification.

**Fig. 7.**
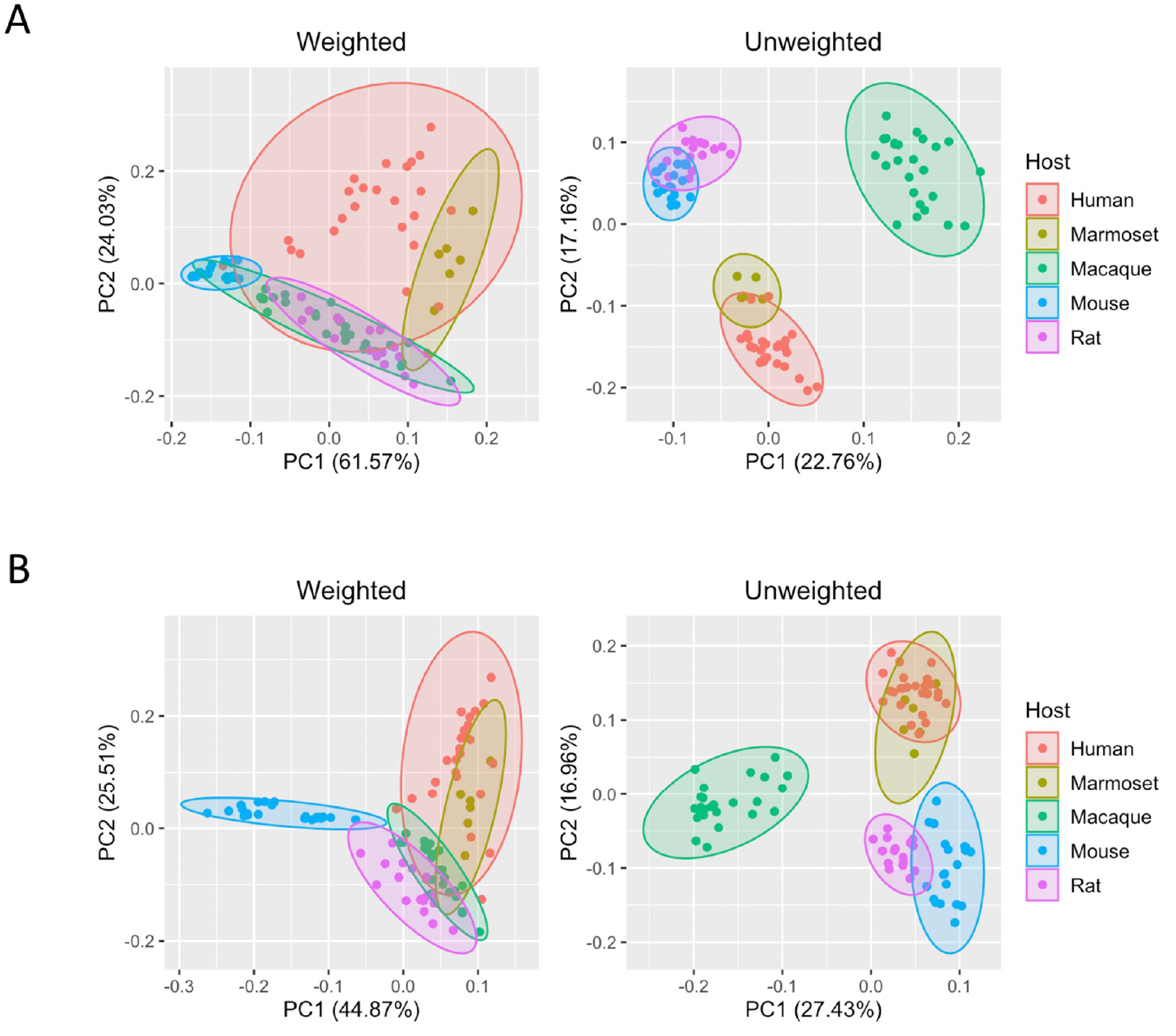
Comparison of the faecal microbiomes of marmosets, mice, rats, macaques, and humans. OTU-based unweighted and weighted PCA (A) at the genus level and (B) at family levels. The 16S rRNA gene sequence data for faecal samples from 6 marmosets were sequenced in this study. The 16S rRNA gene sequence data for faecal samples from humans, macaque monkeys, rats, and mice were obtained from a previous study (34). Weighted (quantitative) accounts for microbiome abundance and unweighted (qualitative) is based on their presence or absence.

### Comparison of microbiomes among animal models by 16S rRNA gene sequencing

To find similarities and differences between the common marmoset microbiome and those from the human and major model animals, macaques, mice and rats, 16S rRNA amplicon sequencing for marmoset faecal samples was conducted. The 16S rRNA gene sequence data for faecal samples from humans, macaque monkeys, rats, and mice were obtained from a previous study (34). The OTU (operational taxonomic unit) analysis of microbiome similarity was performed quantitatively (weighted) and qualitatively (unweighted) at the genus and family levels. The principal component analysis (PCA) of the OTU profiling data is shown in Fig. 6. In contrast to the weighted analysis, the unweighted analysis more clearly isolated clusters of species. The marmoset clusters overlapped with human clusters in both weighted and unweighted analyses, revealing that the marmoset and human microbiomes were most similar. Mouse and rat clusters were located nearby in the unweighted analysis. The analysis at the family level showed that *Muribaculaceae* family accounted for approximately half of the microbiome of mice and was also detected in rat and macaque individuals. In contrast, most of humans and marmosets did not retain *Muribaculaceae* (Fig. S3). Despite all three groups being primates, the macaque microbiome did not resemble marmoset or human microbiomes in the unweighted profile, and no specific bacteria were detected between macaques and humans or between macaques and marmosets. On the other hand, characteristic bacteria were found in the comparison between marmosets and humans. The *Bacteroidaceae* family and *Bacteroides* genus were major members in marmosets and humans. *Bacteroides*, which inhabits healthy human intestines, has been reported to have a reduced abundance in IBD patients and is attracting attention as a probiotic (35). *Bifidobacteriaceae* family, *Bifidobacterium* genus, and *Coriobacteriaceae* family, *Collinsella* genus, were also mostly present in marmosets and humans but were not detected in many individuals of other animal model species. *Bifidobacterium* is known to be significantly depleted in colorectal cancer, IBD, irritable bowel syndrome and obesity, and has been reported to enhance the effectiveness of cancer immunotherapy (36)(37). *Collinsella* is a proinflammatory genus involved in rheumatoid arthritis and non-alcoholic steatohepatitis, and has potential as a disease biomarker (38)(39). In brief, it was found that marmoset and human faecal microbiome are significantly close and share many bacteria involved in a variety of human diseases.

## Discussion

The proposed method reconstructed the common reference metagenome by merging scaffolds from three sites assembled from metagenomic read data; using this approach, it was possible to identify the corresponding genes between three intestinal sites with high accuracy. Here, we evaluated the non-chimeric rate of the reconstructed genomes using a benchmarking dataset that collected only DNA reads assigned to known bacterial species. The non-chimeric rate is defined by the percentage of genome length assembled solely with DNA reads from a single species. As a result, the non-chimeric rates were 92.8% and 94.7% for individual 1 and 2, respectively; most genomes were completely reconstructed as a single species within the metagenome (Supplementary Note 4).

The gene expression changes between the caecum, transverse colon, and faeces were shown to be more dynamic than changes in microbiome abundance, which was consistent with the results of a previous study (5). For example, we found that genes related to carbohydrates were activated in the caecum compared to faeces, and coenzyme metabolism genes and antibacterial resistance genes were more highly expressed in both the caecum and transverse colon than in faeces, but these gene abundance did not vary significantly. Since the differential expressions of these genes were considered to be influenced by the concentration of SCFAs converted from carbohydrates by the microbiome, we focused on the *fucI* gene involved in carbohydrate metabolism. The reconstructed reference metagenome identified 30 bacteria coding *fucI*, and *Megamonas* genus contributed most to the expression of *fucI* at whole community, despite the most abundance of *fucI* of *Akkermansia*. SCFAs are involved in host lipid metabolism (40), and *Akkermansia*, SCFA-producing bacteria, has received attention as a factor that suppress high-fat diet-induced metabolic disorders, including metabolic endotoxemia and insulin resistance (33). Our results show that *Megamonas* is a more important member as a potential producer of SCFAs, especially in the caecal environment. These results highlight that the integrated analysis of metagenomics and metatranscriptomics provide also biological interpretations from two aspects: gene abundance and expression levels.

Finally, we compared the faecal microbiome of six common marmosets with that of humans and the major model animals, macaques, mice and rats by the 16S rRNA gene analysis. The marmoset microbiome was found to be most similar to the human microbiome, with *Bacteroides, Bifidobacterium*, and *Collinsella* shared between them. These results suggest that marmosets can be expected to be a useful animal model in the microbiome studies.

In conclusion, this study developed a method for integrating metagenome and metatranscriptome for the analysis of multiple intestinal sites. This analysis method allows to quantify gene expression levels and analyze gene expression changes among intestinal sites including unknown bacterial genes, which was overlooked with conventional methods. As a result of applying this analysis method to the multiple intestinal sites of the common marmosets, we revealed the changes in the internal environment along the intestinal tract may vary the expression pattern of the microbiome, and moreover this microbial change may mutually affect the environment inside the intestine. These findings highlight the importance of database-independent methods in metatranscriptomic analysis to quantify gene expression in the microbiome.

## Materials and Methods

### Sample collection

Common marmosets were housed at the Central Institute for Experimental Animals (Kawasaki, Japan) with free access to a pellet diet (for monkeys, CREA New World Monkey Diet, CMS-1M; CREA Japan, Tokyo, Japan). Two marmosets were selected in this experiment so that sample volumes from all three sites satisfied the requirements of the experimental protocol. Marmosets were sacrificed with pentobarbital overdose and digestive tract was isolated. The gastrointestinal tract of each animal was excised, and the luminal content of each gastrointestinal tract site was collected and divided into for metagenomic samples and for metatranscriptomic samples. The contents were immediately frozen in liquid nitrogen and stored at −80C. Metatranscriptomic samples were crushed and homogenized in solution D containing guanidinium, which inhibit ribonuclease (41), within one week after dissection to protect against the degradation and stored at −80C. Caecal, transverse colonic and faecal contents of marmosets (individual ID: I6289M and the individual ID: I6027M; Table S7) were used for metagenomic and metatranscriptomic analyses. These three sites were targeted at the beginning, middle and end of the colon, which is abundant in microbiome. Faecal contents of a total of 6 marmosets in addition to these two marmosets were used for 16S rRNA gene analysis.

### Shotgun metagenomic sequencing

DNA was extracted from each metagenomic sample using a MORA-EXTRACT Kit (Kyokuto Pharmaceutical Industrial Co., Ltd., Tokyo, Japan). Sequencing libraries were prepared using TruSeq Nano DNA Library Prep Kit (Illumina Inc, San Diego, CA, USA). All these procedures were performed according to the manufacturer’s instructions (Table S8). Illumina HiSeq sequencing yielded a total of 435 giga nucleotides (Gnt) of paired-end reads (250 bp × 2) for the metagenome. This dataset included an average of 145.1M reads ± 3.9M reads (mean ± s.d.) per sample before quality filtering, described below, and 125.9M reads ± 5.3M reads afterward (Table S9). Shotgun metagenome libraries were adapter trimmed and quality filtered by Trimmomatic (42) version 0.36 (ILLUMINACLIP:Adapter.fa:2:30:10:8:true, LEADING:3, TRAILING:3, SLIDINGWINDOW:4:15, MINLEN:50) and FASTX-Toolkit version 0.0.14 (-q 20 -p 80) (http://hannonlab.cshl.edu/fastx_toolkit/), respectively. Potential host and feed contaminants were then filtered by removing reads with sequences aligned to the host genome and feed genome (Supplementary Note 1).

### Metatranscriptomic sequencing

RNA was extracted by a combination of the acid-guanidium-phenol-chloroform RNA extraction method (43) and a bead crushing method and assessed to ensure high quality (RNA integrity number (RIN) scores ≥ 7.9) (Table S10). The rRNA was removed using a Ribo-Zero Gold rRNA Removal Kit (Epidemiology) (Illumina). Sequencing libraries were prepared using TruSeq Stranded Total RNA HT Kit (Illumina). All these procedures were performed according to the manufacturer’s instructions (Table S8). Illumina HiSeq sequencing yielded a total of 165 Gnt of paired-end reads (100 bp × 2) for the metatranscriptome. This dataset included an average of 137.7M reads ± 2.8M reads (mean ± s.d.) per sample before quality filtering, described below, and 126.7M reads ± 4.0M reads afterward (Table S9). Metatranscriptome libraries were adapter trimmed and quality filtered using the same method as the metagenome libraries. The rRNA reads were removed by SortMeRNA (44) version 2.1 (-e 1e-30). Potential host and feed contaminants were filtered in the same way as the metagenome libraries.

### Integrated metagenomic and metatranscriptomic analyses

The integrated analytical method proposed in this study is composed of three main steps: (i) reconstruction of a reference metagenome common to all sites by assembly, scaffolding and merging steps (Fig.1 (1), (2)), (ii) mapping of DNA and mRNA reads to this reference metagenome respectively (Fig.1 (3), (4)) and (iii) quantification of microbial gene expression levels at whole community and per cell (Fig.1 (5)). Evaluation of this analytical method and determination of parameters for each step were carried out by using genome of known bacterial species genome.

The DNA reads were assembled by Megahit (45) version 1.1.3 (-k-min 21, -k-max 141, -k-step 12, -prune-depth 20). Contigs shorter than 1,000 bp were discarded from further processing. The contigs were scaffolded by OPERA-LG (46) version 2.0.6 using information of paired-end reads information. By merging the scaffolds of metagenomes from three intestinal sites using QuickMerge (47) version 0.3 (-hco 50, -c 50, -ml 1000), the reference metagenomic sequences that were common between sites were reconstructed. Genes were then predicted on the reference metagenomic sequences by MetaGeneMark (48) version 3.38 to make the entire list of genes on the intestinal sites. We used GhostKOALA (49) and DIAMOND blastp (50) version 0.9.21.122 (--evalue 1e-10, --query-cover 85) to annotate the predicted genes according to orthologous groups in the KEGG database (release 94.1) (2) and the COG database (51). Subsequently, mRNA reads was mapped to the metagenomic reference sequences by Bowtie2 (52) version 2.3.4.3 and the number of mRNA reads were counted by HTSeq (53) version 0.9.1 to quantify the gene expression level. DNA reads were also mapped to the metagenomic reference sequences by Bowtie2 version 2.3.4.3 (-x 2000), and the coverage of each metagenomic sequence was calculated by samtools (54) version 0.1.19.

### Co-variation analysis incorporating a bivariate spatial relevance

We performed a co-variation analysis to estimate the function of unknown genes. This analysis is based on the assumption that functionally similar genes are co-variant in their expression levels (23). First, we benchmarked using the profiles of expression at whole community and per cell by assessing the accuracy of this co-variation analysis in classifying the known genes with the same metabolic process. We grouped the known genes into gene clusters by COG annotation and calculated the bivariate spatial association measure (L statistic value) (24) to detect co-varying gene pairs in a six-dimensional vector of expression levels of three sites in two individuals. This benchmark was used to evaluate the model by AUCs and to determine the threshold of L statistic value to guarantee FPR<0.05. As a result of the benchmarking, we found that using the expression levels at whole community were more accurate than using the expression levels per cell. Next, we grouped the unknown genes into gene clusters by protein sequence similarity using MMSEQS2 (55). We used the model determined by benchmarking to perform co-variation analysis on the unknown and known gene clusters together. This allows us to estimate the function of the unknown gene cluster when the known and unknown gene clusters are linked (Supplementary Note 6).

### Quantification of the gene expression level

Metatranscriptomic functional activity was assessed with two manner of quantification methods. The first is a general method to quantify gene expression by normalizing mRNA read counts with transcripts per million (TPM) (this is called “gene expression level at whole community” in this paper). This method can estimate metatranscriptome activity in a microbial community. The second method is to normalize the mRNA read counts with DNA coverage, thus estimating the gene expression level per single bacterium (this parameter is called “gene expression level per cell” in this paper) (Supplementary Note 2).

### Taxonomic profiling

Each reconstructed genome was identified to the taxon level by mapping the predicted genes against the non-redundant protein database and assigning taxonomic annotation with voting based approach using CAT version 4.3.3 (56).

### 16S rRNA gene sequencing and comparison among animal models

To compare the common marmoset faecal microbiomes with those of humans and other major animal models, 16S rRNA sequencing was conducted on the faecal samples from 6 marmosets. Marmoset faecal DNA was extracted from each metagenomic sample using a MORA-EXTRACT Kit (Kyokuto Pharmaceutical Industrial Co., Ltd., Tokyo, Japan) by the bead crushing method. The 16S rRNA V3–V4 amplicon was amplified using a KAPA HiFi HotStart ReadyMix PCR Kit (KAPA BioSystems, USA). the amplicon PCR forward primer (5’-CCTACGGGNGGCWGCAG-3’) and amplicon PCR reverse primer (5’-GACTACHVGGGTATCTAATCC-3’) were used. Sequencing libraries were prepared using a Nextera XT Kit (Illumina) (Table S9). All these procedures were performed according to the manufacturer’s instructions. Sequencing was performed using an Illumina MiSeq sequencer (Illumina) with paired-end reads (forward: 350 bp, reverse: 250 bp). Illumina MiSeq sequencing yielded a total of 11.3 Gnt of paired-end reads (350 bp, 250 bp). This dataset included an average of 3,154K reads ± 1,190K reads per sample before quality filtering and 1,387K reads ± 346K reads afterward (Table S9). The sequences were analysed using QIIME (Quantitative Insights into Microbial Ecology; version 1.9.1) (57). The 16S rRNA gene sequence data for faecal samples from humans, macaque monkeys, rats, and mice were obtained from a previous study (34). To avoid any bias from different sequencing depths, the OTU table was rarefied to the lowest number of sequences per sample.

## Data availability

All raw sequence data have been submitted to the DDBJ under project PSUB014668 from the Ministry of Education, Culture, Sports, Science and Technology of Japan.

## Acknowledgements

Not applicable

## Author s’ Contributions

M.U. performed experiments, conducted bioinformatics analysis and co-wrote the paper; T.I. and E.S. provided samples for the metagenome, metatranscriptome and 16S rRNA gene sequencing; M.U., M.K. and S.H performed DNA extraction and sequencing for the 16S rRNA gene analysis; A.T. performed deep sequencing with high-throughput sequencers for the metagenome and metatranscriptome analysis; Y.S. designed and supervised the research, analysed the data, and co-wrote the paper. All authors have read and approved the manuscript.

## Ethics approval and consent to participate

The animal experiment protocol was approved by the CIEA Institutional Animal Care and Use Committee (approval nos. 17031, 18032 and 19013). The study was conducted in accordance with the guidelines of CIEA that comply with the Guidelines for Proper Conduct of Animal Experiments published by the Science Council of Japan. Animal care was conducted in accordance with the Guide for the Care and Use of Laboratory Animals (Institute for Laboratory Animal Resources, 2011).

## Consent for publication

Not applicable

