## Supplementary Information and Figures for "Development of a metatranscriptomic analysis method for multiple intestinal sites and its application to the common marmoset intestine"

**Supplementary Figure**

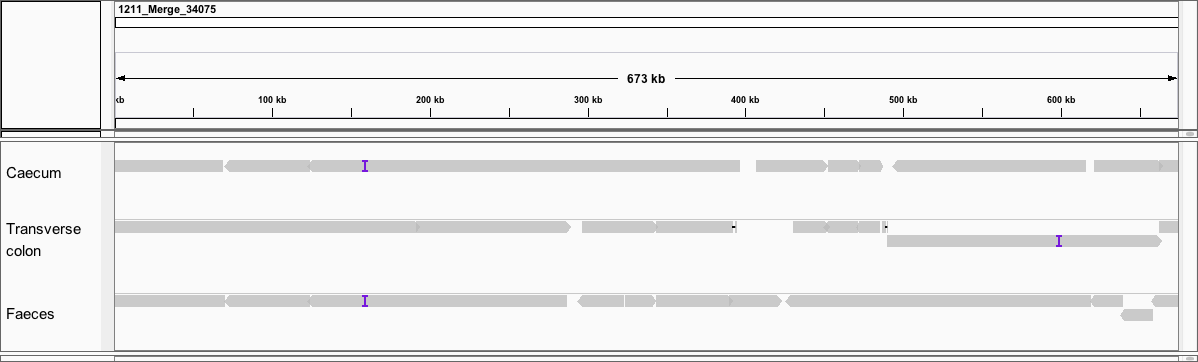

Fig. S1 Visualization of scaffold alignment to the merged genome (MG_34075) by IGV, Related to Fig. 2A

The grey segment represents the scaffold in each site and the purple markers represents an insertion within the scaffold. The scaffolds across the sites complement each other to reconstruct a large genome (Supplementary Note 8).

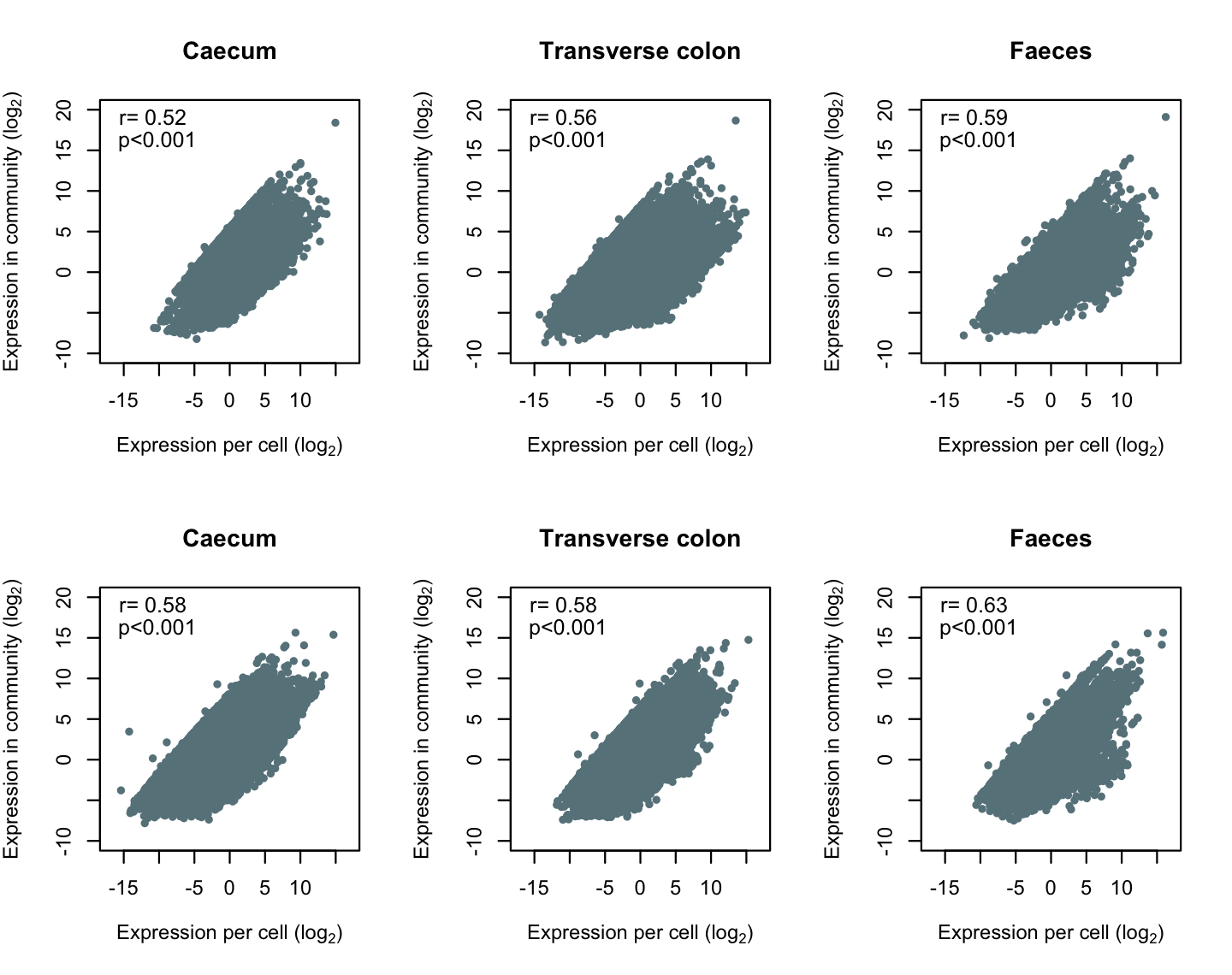

Fig. S2 Pearson correlation coefficient (r) and p-value (p) between the gene expression level at whole community and that per cell

The x-axis indicates the gene expression level per cell, and the y-axis indicates the gene expression level of the entire community. The value converted to log_2_ is used for the expression level. The data points represent expressed genes detected on the reconstructed metagenomes.

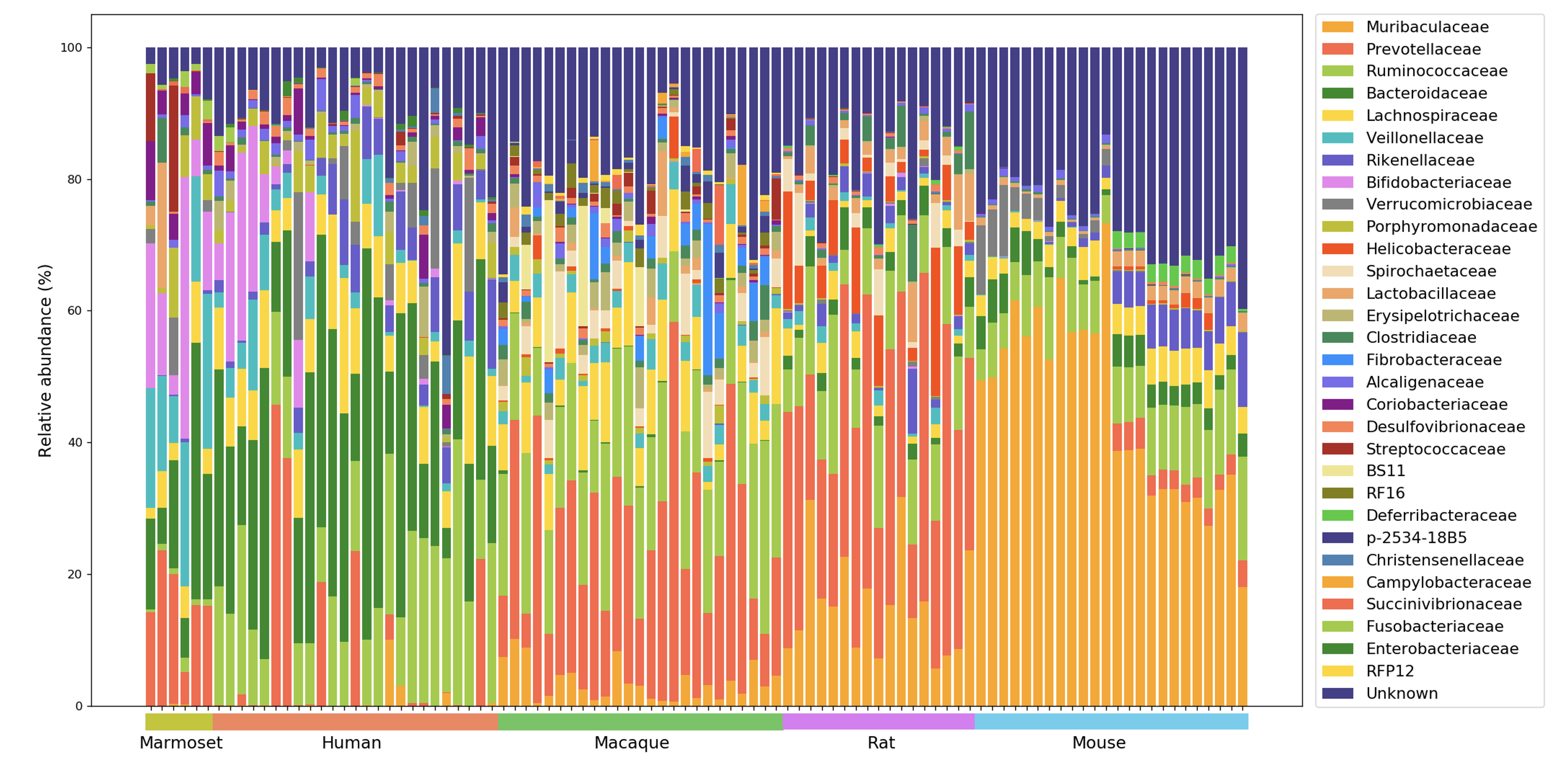

Fig. S3 Relative abundance of bacterial families in marmosets, humans, macaques, rats, and mice, Related to Fig. 7

The 16S rRNA gene sequence data for faecal samples from 6 marmosets were sequenced in this study. The 16S rRNA gene sequence data for faecal samples from humans, macaque monkeys, rats, and mice were obtained from a previous study (1).

**Supplementary Notes**

1. **Removal of contaminated sequences**

To filter the host and feed genomes from the sequenced DNA and RNA reads, these reads were aligned into the common marmoset (Callithrix jacchus; GenBank assembly accession GCA_000004665.1), bread wheat (Triticum aestivum; GenBank assembly accession GCA_900519105.1), soybean (Glycine max; GenBank assembly accession GCA_000004515.4) and (Gadus morhua; GenBank assembly accession GCA_902167405.1) genomes by bowtie2, and the aligned reads were removed.

2. **Computation method of gene expression level per cell**

The functional activity of the microbiome was assessed by the gene expression level at both the whole community and per cell levels. The gene expression levels at whole community were normalized to corresponding gene abundance to obtain an estimate of gene expression levels per cell *T* as follows:

$$T_{g,s,i}=\left\{ \begin{aligned} \frac{R_{g,i}}{\sum_{g} R_{g,i}}\times\frac{\sum_{s\ni g} D_{s\ni g,i}}{D_{s\ni g,i}} , if D_{s\ni g,i}>0 \\ 0, if D_{s\ni g,i}=0 \end{aligned} \right.$$

$R_{g,i}$ is the count of mRNA reads, in TPM, of gene g in sample $i$. $D_{s\ni g,i}$ is the count of DNA reads, in TPM, of the genome sequences where the gene $g$ is located within in sample $i$.

### 3. **Parameter determination and evaluation of the integrated analytical method**

The three steps of the integrated analytical method to reconstruct the metagenome—assembly, scaffolding and merging—were assessed using real DNA reads assigned to the top 20 bacterial species in the taxonomic profile as input (Table S11). For DNA assembly, Megahit was run with all combinations of parameters, as follow: --k-min 35; --k-max 225; --k-step 12; and --prune-depth 2, 5, 10 or 20. For reference metagenomic sequence construction, QuickMerge was run with the following parameters: -hco 5, 10, 50 or 100; -c 1.5, 10, 50 or 100; and -ml 1,000, 5,000 or 10,000.

To assess the accuracy of the genome constructed at each step, the number of genes covered for each reconstructed genome were used as criteria. Upon aligning all reads in the database containing the genomes of the top 20 bacterial species using Blastn (2) (-evalue$<$1e-10, alignment length$\geq$500 bp, identity$\geq$85%), one reconstructed genome aligned to multiple bacterial genomes was defined as a chimaera. The covered genes represent the number of perfectly matched genes upon alignment of the genes against the reconstructed genome. For only the merging step, the number of genes was counted as 1/N if the same gene appeared multiple times (N), which can determine the parameters that were merged correctly. We detected parameter at assembly and merge steps by assessing the number of genes covered by non-chimera genome (Table S12 and S13).

4. **Assessment of reconstructed metagenomics**

We evaluated the integrated analytical method using the dataset constructed from top 20 bacterial species in the taxonomic profile by Kraken2 (Table S11). The DNA reads from the dataset were reconstructed through the assembly, scaffolding and merging steps. The reconstructed genomes were assessed by the non-chimera rate, which is the ratio of the length of all chimeric sequences to the total length of reconstructed genomes. A non-chimaera rate = 1 indicated that the genome was reconstructed with high accuracy without chimera. The chimeric sequences refer to a mixture of genomes from multiple species, which was calculated by aligning the sequences of 20 species in the database to the reconstructed genome using Blastn (-evalue$<$1e-10, alignment length$\geq$500 bp, identity$\geq$85%).

5. **Percentage of genes that match in scaffolds of all 3 sites and the reconstructed scaffold**

To confirmed that the genes were retained before and after the merge, we examined the percentage of genes common to the three sites that match the genes in the reconstructed metagenome. First, we aligned each site scaffold to the reconstructed scaffold for the gene position adjustment. Specifically, we converted the positions of the gene regions of the scaffolds at each site to the positions on the reconstructed scaffold based on the PAF file with cs tags from the alignment result. We then extracted only the genes common to all three scaffolds, excluding genes on scaffolds constructed from one or two sites, to avoid overestimating the percentage of gene matches before and after the merge. We targeted the known genes and calculated the mismatch (including insertin, deletion, variant) of gene regions between the scaffold of the three sites and the reconstructed scaffold (Table S3).

6. **Functional annotation for unknown genes by co-variation analysis**

We performed a co-variation analysis by modifying some of the methods of previous studies (3) to estimate the function of unknown genes. This analysis is based on the assumption that functionally similar genes are co-variant in their expression levels. As shown below, we first benchmarked the expression profiles of the known genes and then estimated the function of the unknown genes using the models determined in this benchmark as follows.

**Benchmarking with known genes**: Benchmarking was performed to determine and evaluate the model that most accurately discriminates genes with common metabolic processes. Known genes are those annotated by the COG database and the KEGG database, as described in "Integrated metagenomic and metatranscriptomic analyses" in this paper. We grouped genes with the same COG ID into known gene clusters and summed the expression levels within each known gene cluster. We excluded gene clusters with an expression variance of less than 1.00 among the 3 sites. The metabolic process of each known gene cluster was applied to the metabolic process of the gene with the longest gene length in the cluster as a representative. In order to relate the genes between individuals, we used the known gene clusters with COG IDs present in both individuals 1 and 2 for co-variation analysis. After these pre-processing steps, we calculated the bivariate spatial association measure (L statistic) (4) of expression levels between all known gene pairs in a six-dimensional vector consisting of three sites in two individuals. L statistic for gene expression levels X and Y was calculated by:

$$L_{X,Y}=\frac{\sum_{i} \left[ \left( \sum_{j} w_{ij}\left( x_{j}-\bar{x} \right) \right)\cdot\left( \sum_{j} w_{ij}\left( y_{j}-\bar{y} \right) \right) \right]}{\sqrt{\sum_{i} \left( x_{i}-\bar{x} \right)^{2}}\sqrt{\sum_{i} \left( y_{i}-\bar{y} \right)^{2}}}$$

where $w_{ij}$is a row-standardized version of a spatial wight $v_{ij}$, which is defined as:

$$v_{ij}=\left\{ \begin{aligned} {d_{ij}}^{-b} (i\neq j) \\ 1 (i=j) \end{aligned} \right.$$

where $d_{ij}$ refers to the distance between site $i$ and site $j$ (Table S14), and $b$ to the distance friction coefficient ($b=2$).

By applying $L$ to covariate analysis, we can calculate the covariation value between two gene expression levels, taking into account the spatial information of the intestinal sites; for example, the distance between the caecum and the transverse colon is closer than the distance between the caecum and the anus (faeces). The known gene cluster pair was defined as a covariate gene if $L$ is greater than a threshold value. True positives (TPs) were defined as pairs of covariant genes with a common metabolic process definition. False positives (FPs) were defined as pairs of covariant genes without common metabolic linkage definition. True negatives (TNs) were defined as pairs of non-covariant genes without a common metabolic process. False negatives (FNs) were defined as pairs of non-covariant genes with a common metabolic process. The ROC curve was plotted by calculating the false positive rate (FPR; $FPR= FP / (FP + TN)$) and sensitivity ($sensitivity= TP / (TP + FN)$) while varying the threshold of the $L$ from 0.00 to 1.00 in 0.01 increments. We assessed the prediction accuracy by calculating the AUC (Fig. 3 A, B and C). As a result of this benchmarking, the model with the highest accuracy (using gene expression profiles at whole community and KEGG reaction as the metabolic linkage definition) was used for subsequent co-variation analysis to estimate the function of unknown genes (Table S5). threshold of 0.885 was applied as a value to guarantee FPR<0.05.

**Estimation of unknown gene functions**: We grouped unknown genes by protein sequence similarity using MMSEQS2 (–cov-mode 1, –cluster-mode 2, -c 0.9, -s 7, –kmer-per-seq 20) (5) as unknown gene clusters and summed the expression levels within each unknown gene cluster. As in the benchmark pre-processing, we excluded gene clusters with a variance of less than 1.00 among the 3 sites and used gene clusters present in both individuals 1 and 2 for co-variation analysis. Using the model determined by the benchmark, the unknown and known gene clusters were combined for the co-variation analysis. We estimated the metabolic process of the known gene cluster as that of the unknown gene cluster when a known gene cluster was linked to an unknown gene cluster. We identified 3,528 unknown gene clusters that are significantly linked to known gene clusters (Table S6). To find potential functional trends of unknown gene clusters, we examined the functional categories that shared by a large number of gene clusters by enrichment analysis with Fisher’s exact test. A functional category was considered significantly different between the known gene clusters and unknown gene clusters if the Benjamini-Hochberg adjusted P value was <0.01 (Fig. 3D).

7. **Validation of co-variation analysis by sequence similarity of linked gene clusters**

In the co-variation analysis, we estimated the function of unknown genes by linking gene clusters using the variation of gene expression levels. We sought to demonstrate that the functions estimated by this co-variation analysis were functions that could not be annotated by sequence similarity. Therefore, we identified linked gene cluster pairs with sequences that are even remotely similar using DIAMOND blastp version 0.9.21.122 (--evalue 0.1) (6), and found that no sequences were similar between the clusters. This result shows that co-variation analysis using gene expression levels links a potential function that cannot be annotated by sequence similarity.

8. **Visualization of scaffold alignment to the merged genome by IGV**

We reconstructed bacterial genome by merging scaffolds among sites. To visualize the scaffolds that construct the merged genome, the scaffolds of each site were aligned with the merged genome by minimap2 version 2.17-r941 (-g 100 -r 100 --no-long-join). A BAM file was loaded to IGV version 2.7.2. The scaffolds across the sites complemented each other to reconstruct a large genome, and the merging improved the contiguity of the genome (Fig. S1).
